# Bayesian Markov models consistently outperform PWMs at predicting motifs in nucleotide sequences

**DOI:** 10.1101/047647

**Authors:** Matthias Siebert, Johannes Söding

**Affiliations:** Quantitative and Computational Biology, Max Planck Institute for Biophysical Chemistry, Am Fassberg 11, 37077 Göttingen, Germany; Gene Center, Ludwig-Maximilians-Universität München, Feodor-Lynen-Strasse 25, 81377,Munich, Germany

## Abstract

Position weight matrices (PWMs) are the standard model for DNA and RNA regulatory motifs. In PWMs nucleotide probabilities are independent of nucleotides at other positions. Models that account for dependencies need many parameters and are prone to overfitting. We have developed a Bayesian approach for motif discovery using Markov models in which conditional probabilities of order *k*-1 act as priors for those of order *k*. This Bayesian Markov model (BMM) training automatically adapts model complexity to the amount of available data. We also derive an EM algorithm for de-novo discovery of enriched motifs. For transcription factor binding, BMMs achieve significantly (*p*<0.063) higher cross-validated partial AUC than PWMs in 97% of 446 ChIP-seq ENCODE datasets and improve performance by 36% on average. BMMs also learn complex multipartite motifs, improving predictions of transcription start sites, polyadenylation sites, bacterial pause sites, and RNA binding sites by 26%-101%. BMMs never performed worse than PWMs. These robust improvements argue in favour of generally replacing PWMs by BMMs. The <u>Ba</u>yesian <u>M</u>arkov <u>M</u>odel <u>motif</u> discovery software BaMM!motif is available under GPL at http://github.com/soedinglab/BaMMmotif.

## Background

The control of gene expression allows the cell to adapt its protein and RNA inventory in response to developmental and environmental cues. At its center lies the binding of proteins to specific motifs in promoters and enhancers to control RNA synthesis rates and to RNAs to regulate their splicing, localisation, translation and degradation. The accurate prediction of protein binding affinities to DNA and RNA sequences is therefore of central importance for a quantitative understanding of cellular regulation and of life in general.

Most known models that describe the sequence specificity of transcription factors were deduced from in-vivo binding sites measured by ChIP-seq [1], from in-vitro measurements of binding strengths using either protein binding microarrays (PBMs) [2] or in-vitro selection coupled to high-throughput sequencing (HT-SELEX) [3], or from bacterial one-hybrid assays [4]. To obtain a statistical model of binding specificity from such measurements, motif discovery algorithms learn the model parameters that agree best with the measurements. At present, the standard model for this purpose is the position weight matrix (PWM), and thousands of PWMs for transcription factors are available in motif databases such as JASPAR, HOCOMOCO, SwissRegulon, or TRANSFAC [5, 6, 7, 8].

PWMs rank binding sites according to the log-odds score 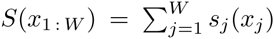 with contributions 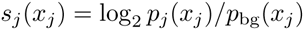 that depend only on single nucleotides *x_j_* in the binding site sequence 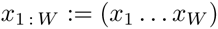. Here, 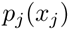 is the probability of nucleotide 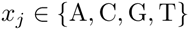 to occur at position *j* of the binding site, and 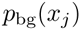 is the background probability for nucleotide *x_j_* in a representative sequence set.

PWMs cannot model correlations between nucleotides. For example, if 50% of binding site sequences are GATC and the other 50% are GTAC, a PWM will give the same high score to GTTC and GAAC as to the true binding sequences. It cannot learn that at position 2 an A must be followed by a T and T by A.

Nucleotide correlations can originate from (1) stacking interactions that determine binding through DNA ”shape readout” [9], (2) amino acids that contact multiple bases simultaneously [10], (3) multiple sequence-dependent binding modes of a factor [11, 12, 13, 14], and (4) complex multi-submotif architectures with varying submotif spacings, which are typically bound cooperatively by multiple factors [15, 16, 17].

For these reasons one might expect more complex models that do not assume nucleotides to contribute independently to the binding strength to perform better than PWMs. However, the usefulness of such more complex models has been controversially discussed for long [18, 19, 20]. Zhao et al. found PWMs to be as accurate as mixtures of PWMs to describe the binding strengths of transcription factors measured by PBMs [21, 2]. Weirauch et al. concluded that for >90% of tested transcription factors PWMs performed as well as more complex models to predict PBM binding strengths, and they were just as good in predicting in-vivo ChIP-seq binding sites for all factors [22]. In a recent study, 1’st-order Markov models performed significantly better than PWMs in predicting in-vivo binding sites for 21 % of 96 tested datasets [5]. It is still not clear whether these results are really due to the absence of correlations in binding sites of most transcription factors or to what extent they are explained by the difficulty to train the many model parameters reliably and robustly.

Numerous models incorporating nucleotide dependencies have been developed to improve the modelling of binding site motifs and complex, multipartite motifs. Some learn mixtures of PWMs [23, 24, 25] or Markov models [26], or profile hidden Markov models (HMMs) [27]. But dependencies generally decrease with increasing distance [3], and therefore most models are based on inhomogeneous Markov models (iMMs), in which the probability of *x_j_* depends on the previous *k* nucleotides *x_j-k:j-1_* [28, 29, 30].

The drawback of iMMs is that the number of parameters *W* x 3 x 4*^k^* grows exponentially with *k*. Already for a 2’nd-order model we need 48 parameters per position. To estimate them with 10% accuracy requires ~100 counts per 3-mer, or 4 800 sequences. When fewer sequences are available, more complex models risk being overtrained: they may perform significantly worse than a simple PWM model due to the noisy parameter estimates while showing overly optimistic performance on the training data.

To prevent overtraining, various heuristic methods were suggested that reduce the number of parameters in a data-driven fashion, by pruning the dependency graph describing which positions each motif position *j* depends on [23, 31, 32, 33]. These methods have several technical drawbacks: (1) they take yes/no decisions, which necessarily lead to a loss of information near the decision boundary. (2) Optimising a discrete dependency graph is cumbersome: to decide between two alternative graphs one needs to find the optimum model parameters for each graph. Also, since two graphs usually induce models with different numbers of parameters, a likelihood-based optimisation is not possible. (3) The discreteness of the graph topology precludes efficient gradient-based optimisation techniques. (4) Finally, no algorithms have been put forward to train these models on unaligned motifs. These models can therefore not be applied for de-novo motif discovery.

Here, we present a Bayesian approach to learn inhomogeneous Markov models for sequence motifs that makes optimal use of the available information while avoiding overtraining. The key idea is that we use the conditional probabilities of order *k* - 1 as priors for the conditional probabilities of order *k*.

Our Bayesian approach is similar to interpolated Markov models [34, 35] in that the probabilities of order *k* are obtained as linear interpolation of the maximum likelihood (ML) estimate for order *k* and the lower-order probabilities. Various rather ad-hoc methods have been used to set the interpolation weights [34, 35, 36, 37] (e.g. by making them depend on the p-value with which the hypothesis can be rejected that the conditional probabilities of order *k* - 1 and of order *k* are noisy estimates of the same underlying distributions [36]). In contrast, the interpolation weights of BMMs emerge naturally from our probabilistic approach without further assumptions except for the choice of priors.

We analyse how much can be gained by using higher-order inhomogeneous BMMs over two baseline methods: 0’th-order BMMs, which are simply PWMs trained with the standard EM-type algorithm as implemented in MEME [38], and our tool XXmotif, which performed favourably in comparison to state-of-the-art motif discovery tools [39]. We assessed these different methods by the quality of the models they produced starting from the motif occurrences discovered by XXmotif. We demonstrate consistent improvements by higher-order BMMs as compared to PWMs on each of a large and heterogeneous collection of datasets with simple and complex motif architectures. Likewise, correlation of predicted binding affinities with quantitative EMSA measurements was substantially improved.

## Methods

### Bayesian Markov model learning

We would like to solve the following task: We have a set of sequences that are enriched in a sought motif, for example a binding motif for a transcription factor or a multi-submotif region with complex architecture. The sequences might have been produced by ChIP-seq or SELEX-seq experiments, they might be promoters of coregulated genes, regions of differential DNase I accessibility, or specific genomic sites such as splice sites. Our goal is to train a model that discovers the location and strength of enriched motifs in the training sequences and that can predict motifs and their strengths in arbitrary sequences.

In the Supplemental Methods (eqs. S.1-S.7) we show that, to learn a model for the Gibbs binding energy 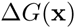 to sequence 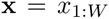, we can solve an equivalent statistical learning problem: we learn the probability distribution of motif sequences, 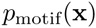, and of background sequences, 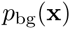. Then the binding energy in units of *k*_B_*T* (where *k*_B_ is Boltzmann’s constant) is, up to a constant, given by the log-odds score, 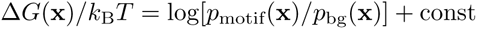. Here we rely on the common approximation of unsaturated binding (low factor concentration), whereby the probability to observe sequence x in the training set is proportional to 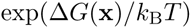.

We model the background sequences with a homogeneous Markov model (MM) of order *K’*, 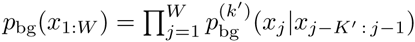 and the motif sites with an inhomogeneous Markov model (iMM) of order *K*, 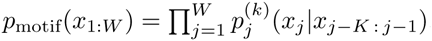. This results in a log-odds score

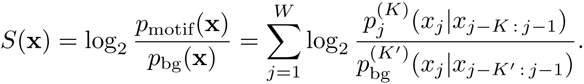

Since the binding energy 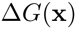 is a linear function of the log-odds score, the score is ideal for ranking potential binding sites by their predicted strength.

The central idea for Bayesian Markov Model (BMM) training is that, to learn the *k*’th-order probability 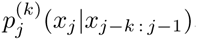, we can use the order-(*k*-1) probability 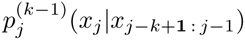 as prior information. The latter is an excellent approximation of the former because dependencies between positions generally decrease quickly with increasing distance [3]. And since the shorter context 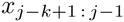 is on average four times more frequent than 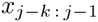, the lower-order probabilities will also be more robustly estimated.

We learn the parameters of the inhomogeneous Markov model by maximising the posterior probability the product of the likelihood and the prior probability (eq. (S.12) in Supplementary Methods). A natural prior is a product of Dirichlet distributions with pseudocount parameters proportional to the lower-order model probabilities, with proportionality constants *α_k_* for *k =* 1,…, *K* whose size determines the strength of the prior. Maximizing the posterior probability yields

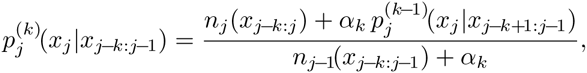

which is illustrated in Figure 1. For frequently occurring *k*-mers 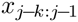 the counts dominate over the pseudocounts and we can accurately estimate the conditional probabilities from the counts. For *k*-mers with few counts the pseudocounts dominate and the probability reverts to the estimate at order *k -* 1, which in turn may be dominated by the estimate at order *k -* 2, and so forth, down to an order where the number of counts dominates the pseudocounts. In this way, conditional probabilities are learned only for those k-mers for which they can be robustly estimated, while other conditional probabilities are approximated by robustly estimated lower-order probabilities.

**Figure 1.**
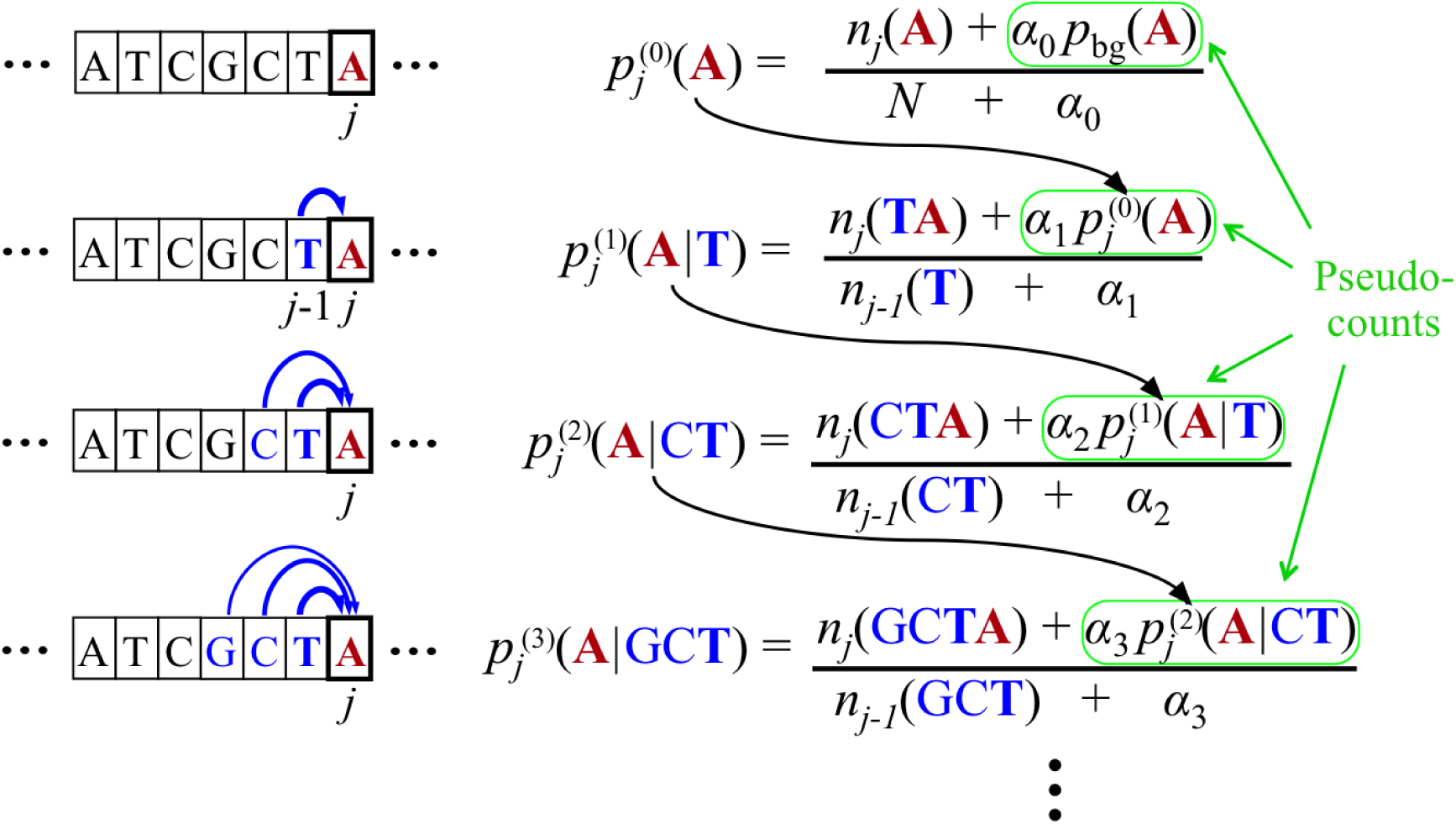
**Bayesian Markov model training automatically adapts the effective number of parameters to the amount of data**. In the last line, if the context GCT is so frequent at position *j* in the motif that its number of occurrences outweighs the pseudocount strength, 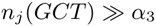, the 3’rd-order probabilities for this context will be roughly the maximum likelihood estimate, e.g. 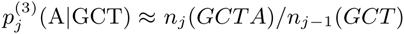. However, if few GCT were observed in comparison with the pseudocounts, 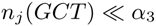, the 3’rd-order probabilities will fall back on the 2’nd-order estimate, 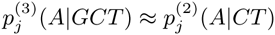. If also 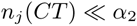, then likewise the 2’nd-order estimate will fall back on the 1’st-order estimate, and hence 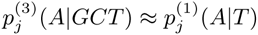. In this way, higher-order dependencies are only learned for the fraction of *k*-mer contexts that occur sufficiently often at one position *j* in the motifs training instances to trump the pseudocounts. Throughout this work we set 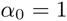 and 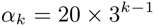.

We fixed the prior strengths 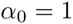 and 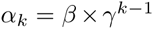 for 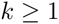, with hyperparameters 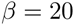 and 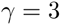. The increasing strength of pseudocounts with increasing *k* reflects the prior belief that dependencies should quickly decline with distance [3]. Owing to this rather strong regularisation, we prevent overtraining on all datasets (see, e.g. Supplementary Figure S18). As background model, we always train a 2’nd-order homogeneous BMM on the set of positive training sequences, with the default *α* setting from our tool XXmotif [39] (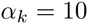 for all 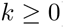). This choice gives good performance as trimers capture the properties of background sequences in sufficient detail without learning the motifs themselves.

### Motif discovery using Bayesian models

When the motif sites in the training sequences are not known a priori, we need to learn a good model and at the same time find motif instances in sequences that are often hundreds or thousands of nucleotides long. Most motif discovery algorithms train PWMs. We derieve here an expectation maximisation (EM) algorithm to train BMMs. A formal derivation is given in the Supplementary Methods.

The goal is to estimate the model parameters 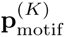, which is a vector containing the 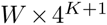 conditional probabilities 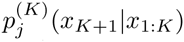 for any 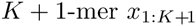. The EM algorithm cycles between E- and M-step. In the E-step, we re-estimate the probabilities for a motif to be present at position *i* of sequence *n*,

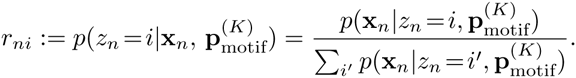

We use the zero-or-one-occurrence-per-sequence (ZOOPS) model [38], and the hidden variable 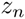 indicates at which position the motif is present in sequence *n*. In the M-step we use the new 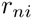 to update the model parameters 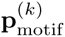 for all orders *k =* 0,…, *K*. This update equation looks exactly the same as the previous equation for known motifs locations, except that now the counts 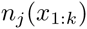 are interpreted as fractional counts computed according to

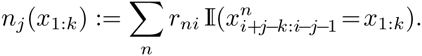

The update of model parameters in the M-step runs through all orders from *k =* 0 to *k = K*, each time using the just updated model parameters from the order below. We iterate the EM algorithm until convergence.

## Results

### Nucleotide dependencies in transcription factor binding sites

We show how BMMs can improve by 29 % the prediction of transcription factor binding sites learned from ChIP-seq-enriched sequences by modeling the correlations between nucleotides in their binding sites.

We evaluated our approach on 446 human ChIP-seq datasets for 94 sequence-specific transcription factors associated with RNA polymerase (RNAP) II from The ENCODE Project Consortium [1]. Positive sequences were compiled from up to 5 000 peak regions with highest confidence by extracting ±102 bp around peaks. We initialised our BMM learning with the motif instances that XXmotif uses to build its top PWM model but added two positions to both 5′- and 3′-ends of the models. We sampled background sequences with the same length as the positive sequences but 100 times as many, using the trimer frequencies from the positive sequences. To compare PWMs with higher-order BMMs without any influence from other source except model order, we treated PWMs as 0’th-order BMMs.

The performance in discriminating binding from background sequences was assessed using four-fold cross-validation, by sorting in descending order the sequences by their maximum log-odds score over all possible motif positions and recording the cumulated number of correct predictions (TP) and false predictions (FP) above a score threshold. Lowering the score threshold from maximum to minimum we trace out a curve of TP versus FP. The normalised version in which one plots the true positive rate (TPR), the fraction of TP out of all positive sequences, versus the false positive rate (FPR), the fraction of FP out of all background sequences, is called receiver operating characteristic (ROC) curve. The partial area under the ROC curve (pAUC) up to the 5’th percentile of FPR (insets in Figure 2A,B) is a good measure of performance, because at FPR > 0.05 and TPR = 1 the precision, i.e. the fraction of predictions that are correct (i.e. truly bound), has already fallen to below 1/(1 + 0.05 × 100) = 0.167. The pAUC therefore summarises the part of the ROC curve most relevant in practice for predicting factor binding sites and is preferable over the AUC.

**Figure 2.**
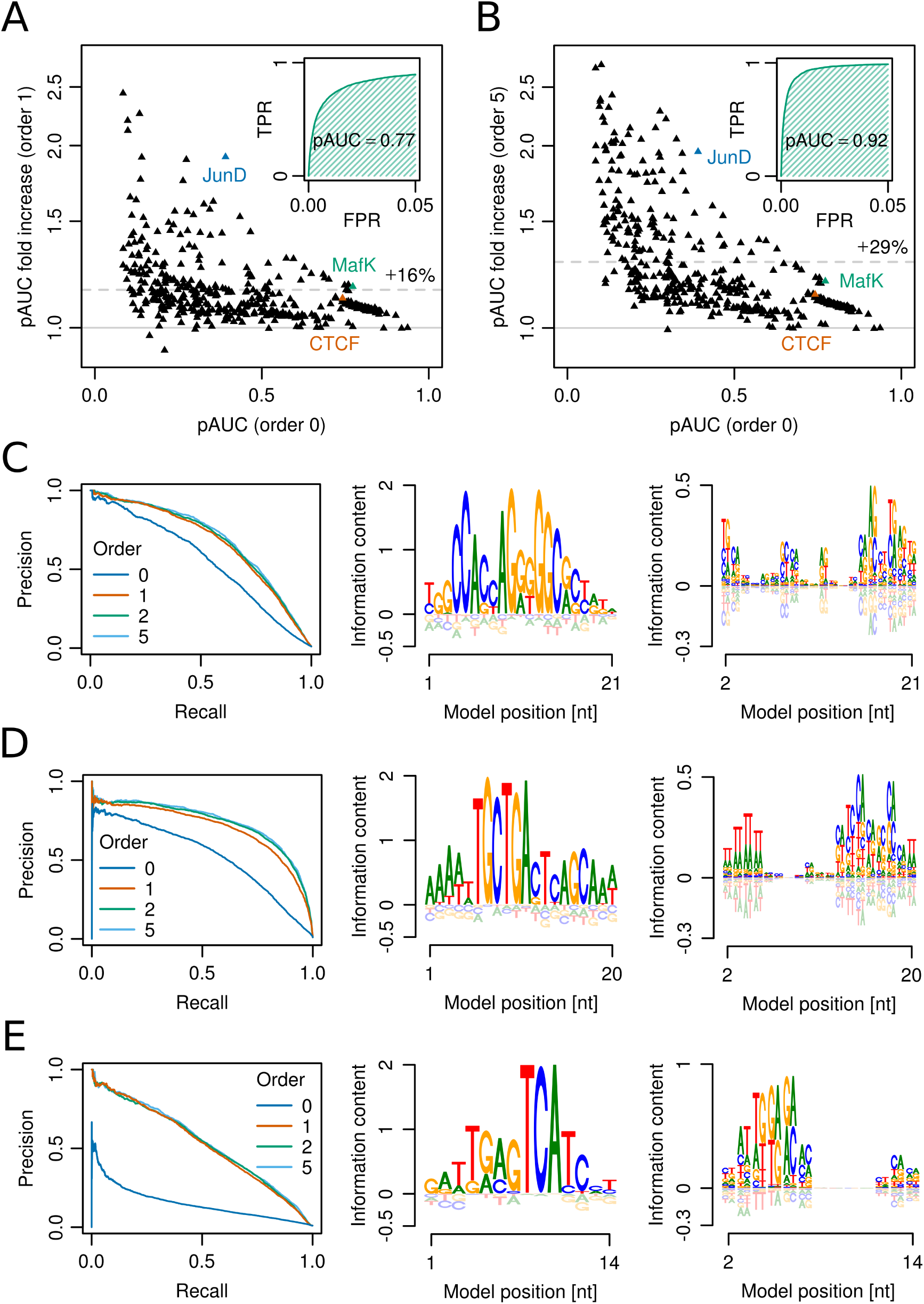
**Modelling nucleotide dependencies in transcription factor binding motifs improves motif discovery and prediction. (A)** Factor of increase in partial area under the ROC curve (pAUC) of 1’st-order BMMs versus 0’th-order BMMs (PWMs) on 446 ChIP-seq datasets for transcription factors from ENCODE. The average performance increase is 16% (dashed line). Y-scale is logarithmic. Inset: partial ROC curve for PWM of MafK binding. **(B)** Same as A but showing the increase in pAUC of 5’th-order BMMs versus PWMs. Inset: 5’th-order BMM of MafK binding. **(C)** CTCF models learned from ChIP-seq sites in Mcf-7 cells. Predictive performance (left) for BMMs of increasing order. 0’th-order (middle) and 1’st-order (right) sequence logos of 2’nd-order BMM. **(D)** Same as C for MafK binding measured in HepG2 cells. **(E)** Same as C but for JunD binding in HepG2 cells.

Figure 2A shows the ratios of pAUC for 1’st-order BMMs to the pAUC for PWMs (order 0) on each of the 446 datasets. On almost all sets the pAUC increases and the average relative increase is 16%. Strikingly, 5’th-order BMMs perform considerably better than 1’st-order BMMs, yielding an average fold pAUC increase over PWMs of 29 % (Figure 2B). Also, on none of the 446 dataset they are clearly worse than PWMS, showing that overtraining is effectively prevented. Higher-orders are particularly beneficial for the more challenging datasets with low pAUC values.

Figure 2C-E illustrates the improvements for specific datasets. The precision-recall curves summarise predictive performance, showing the precision TP/(TP + FP) versus the recall (= sensitivity), the fraction of all bound sequences that are predicted at this precision.

We developed *sequence logos for higher orders* to visualise the BMMs. We split the relative entropy 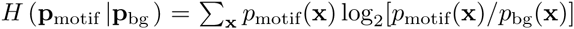 into a sum of terms, one for each order. The logos show the amount of information contributed by each order *over and above what is provided by lower orders*, for each oligonucleotide and position.

The well-studied CCCTC-binding factor (CTCF) has been implicated in the establishment of topologically associating domains and the formation of regulatory chromosome interactions [40]. A 5’th-order BMM for CTCF achieves 14% higher pAUC than a PWM (orange triangle in Figure 2B, Figure 2C, left). The first-order sequence logo identifies the added information (Figure 2C, right). For example at position 16 an A is preferentially followed by a G and a G by a C, relative to the 0’th-order model. The 1’st-order dependencies may reflect the intricate interplay of a subset of CTCF’s 11 zinc-finger (ZnF) domains.

Transcription factor MafK of the AP-1 family of basic-region leucine zippers (bZIP) can bind DNA as homodimer or heterodimer. Depending on its multimeric state, MafK targets the 13 bp T-MARE or the 14 bp C-MARE motif. These are composed of a 7bp and 8bp core sequence, respectively, flanked by GC elements on both sides [41]. A 5’th-order BMM for MafK achieves a 19% higher pAUC than a PWM (green triangle in Figure 2B). Most of this improvement is already present in 1’st order (Figure 2D, left). The 1’st-order logo shows that two alternative DNA recognition modes are represented by the BMM. While the T-MARE is primarily modeled in 0’th order (Figure 2D, middle), the C-MARE is modeled via 1’st-order dependencies (Figure 2D, right). The upstream AT-rich region seen in 0’th order is revealed by the 1’st-order logo to be a poly(dA:dT) tract. This indicates that MafK reads out the narrowed DNA minor groove width known to be induced by poly(dA:dT) tracts [42].

The bZIP transcription factor JunD binds two half-site motifs separated by one or two base pairs. The preferred spacing is cell-type-specific and depends on the availability of oligomerization partners [43]. A 5’th-order BMM for JunD achieves a 96 % higher pAUC than a PWM (blue triangle in Figure 2B, Figure 2E left). The 0’th-order model represents the two aligned right half-sites and a mixture of the two left half-sites displaced from each other by 1 bp (Figure 2E middle). The 1’st-order model deconvolutes the two overlaid half-sites, encoding the sequences ATGAC and XATGA at positions 2–6 (Figure 2E, right).

Supplementary Figures S1-S9 contain analyses for further transcription factors: BATF, c-Jun, c-Fos, Hnf4a, IRF4, NF-YB, NRSF, PU.1, and ZnF143. Remarkably, for some datasets, e.g. ZnF143, we still observe substantial improvements at order three or higher.

### Improvements from flanking nucleotides

We show here that, by including the nucleotides flanking the core binding sites of transcription factors, we can substantially increase the predictive performance of BMMs but less so for PWMs, widening the performance gain of BMMs over PWMs to 36%.

Two recent studies pointed out that for some transcription factors the nucleotides flanking the core binding site make sizeable contributions to the binding specificity [44, 45]. These contributions are probably owed to shape readout around to the core site, for example the minor groove width [42]. Since DNA shape is largely determined by base-stacking interactions, we expect shape readout to lead to strong next-neighbour nucleotide dependencies. If this is true, higher-order models should profit particularly from the inclusion of flanking nucleotides.

We therefore analysed the contribution of flanking nucleotides on BMMs of various orders by comparing models of the same length as found by XXmotif with models extended by four base pairs on either side. For the 8-bp-extended models, we also extended all sequences by 8 bp to keep the same search space size. We then trained original and 8-bp-extended BMMs.

PWMs increase their pAUC by 5% on average (Figure 3A), with a few datasets showing considerably better and others considerably worse performance with 8-bp-extended models. Nicely, 5’th-order BMMs indeed increased their pAUC much more, by 19% on average, and only a single dataset shows > 5% worse performance with an extended model (Figure 3B).

**Figure 3.**
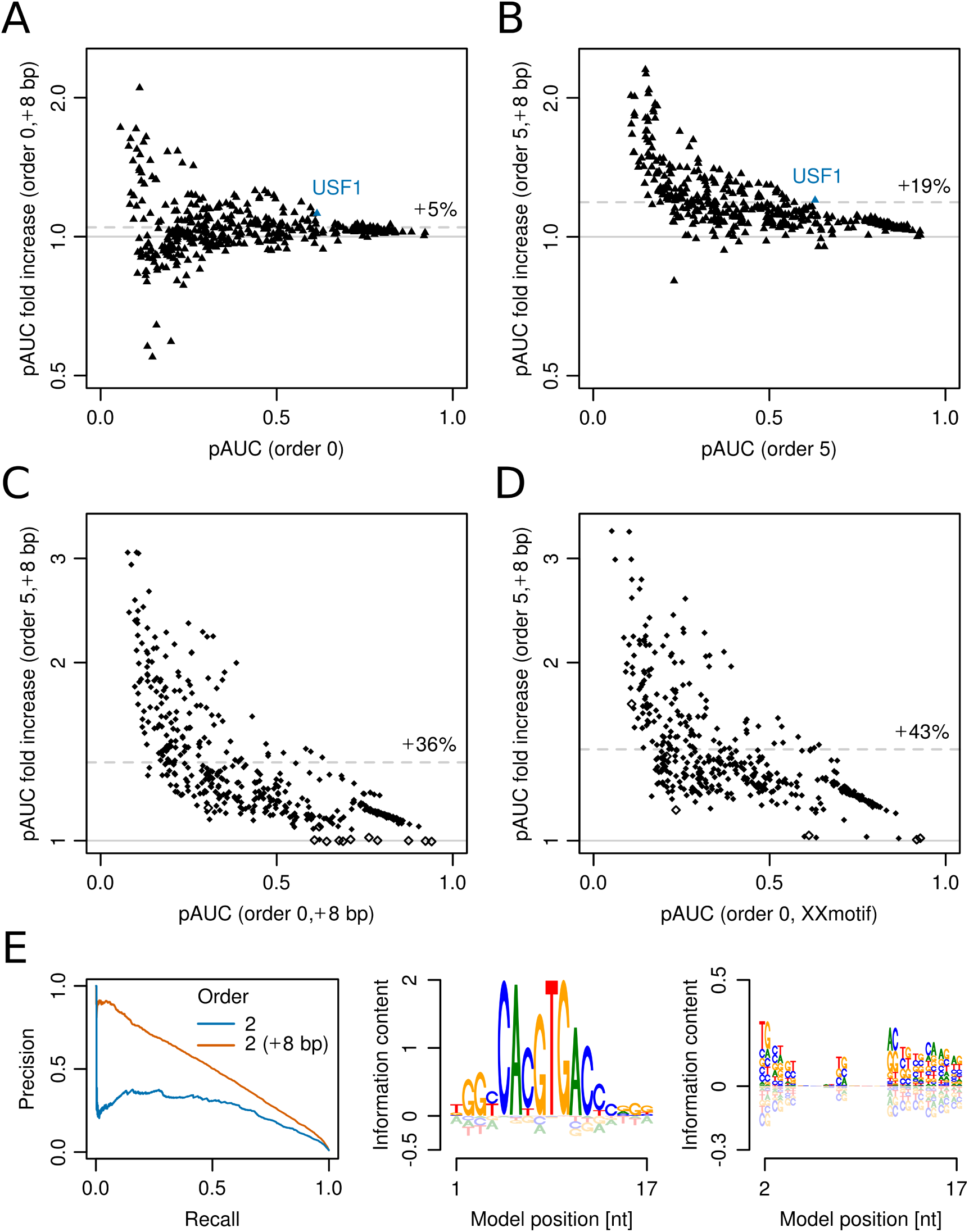
**Nucleotides flanking the core binding sites of transcription factors may contribute greatly to the specificity of higher-order models. (A)** Factor of increase in performance (on log scale) of 8-bp-extended versus unextended 0’th-order BMMs (PWMs) on 446 ChIP-seq datasets for transcription factors from ENCODE. The mean increase is 5% (dashed line). **(B)** Performance increase of 5’th-order 8-bp-extended versus unextended BMMs **(C)** Performance increase of 5’th-order 8-bp-extended BMMs versus 0’th-order 8-bp-extended BMMs (PWMs). Significant improvements (*p* = 6.25%) are obtained on 97% of all datasets (filled diamonds). The remaining 12 datasets show insignificant differences (open diamonds). **(D)** Performance increase of 5’th-order 8-bp-extended BMMs versus PWMs refined by XXmotif. **(E)** Results for 8-bp-extended USF1 model learned from ChIP-seq sites in the H1-hESC line.

When comparing 8-bp-extended PWMs with 8-bp-extended BMMs of 5’th order (Figure 3C), the average increase in pAUC was 36 %. Most strikingly, extended 5’th-order BMMs significantly (at 6.25% level) outperformed the 0’th-order BMMs and also the XXmotif models (Figure 3D) in the vast majority of datasets (97% and 99%, respectively), even though XXmotif compared favourably to state-of-the-art motif discovery tools [39].

Figure 3E shows the results for the 2’nd-order BMM of basic helix-loop helix (bHLH) family transcription factor USF1. The additional information in the flanking regions in 1’st order (right logo) leads to an increase in pAUC of 20 % (blue triangle in Figure 3B) and an increase of precision from 25 % to 90 % for low recall (Figure 3E left). The strong influence of flanking nucleotides has also been demonstrated for other bHLH transcription factors [44, 46], including CBF1, a homolog of USF1 in *S. cerevisiae*. Similar analyses for the transcription factors GR, IRF1, and c-Fos can be found in Supplementary Figure S10.

Piqued by this success, we asked how much can be gained by including a still larger sequence context around core sites. We chose CTCF for its importance in chromatin organisation and extended the core model by 25 bp on either side. Again, only higher-order models profit markedly (Supplementary Figure S11). The predictive performance for the 2’nd-order model reaches an impressive recall of 52% at 95% precision, whereas the 0’th-order BMM predicts only 14% true sites at that precision.

### Quantitative prediction of binding affinities

To attain a quantitative understanding of transcriptional regulation, we need to predict accurately the factor occupancies on regulatory sequences. We demonstrate here that BMMs trained on ChIP-seq data can predict binding affinities directly measured by biophysical methods considerably more accurately than PWMs and a number of competing methods.

Sun *et al*. [47] measured dissociation constants (*K_d_*) of the pioneer ZnF transcription factor Klf4 for binding to various sequences by competitive EMSA experiments. 33 sequences had a single mutation and 25 sequences had multiple mutations to the 10bp Klf4 consensus motif. As in Sun *et al*. [47], we computed the logarithms of each *K_d_* divided by a 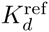. For the sequences with single mutations 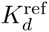 was their median *K_d_* and for the sequences with multiple mutations the *K_d_* closest to their mean.

We trained BMMs of increasing complexity using 101bp sequences extracted around the 5 000 strongest ChIP-seq peaks from [48]. We plotted the log ratios of EMSA *K_d_*’s versus the corresponding predictions from our models. We compared the performance of BMMs of increasing order by means of the Pearson correlation between measured and predicted log *K_d_* ratios.

Overall, the Pearson correlation improves with increasing BMM order (solid lines in Figure 4A, left). While the 0’th-order BMM successfully predicts Klf4 affinities to singly mutated binding sites, it fails for the multiply mutated binding sites (Figure 4A, middle). In contrast, the 5’th-order BMM succeeds on both sets (Figure 4A, right).

**Figure 4.**
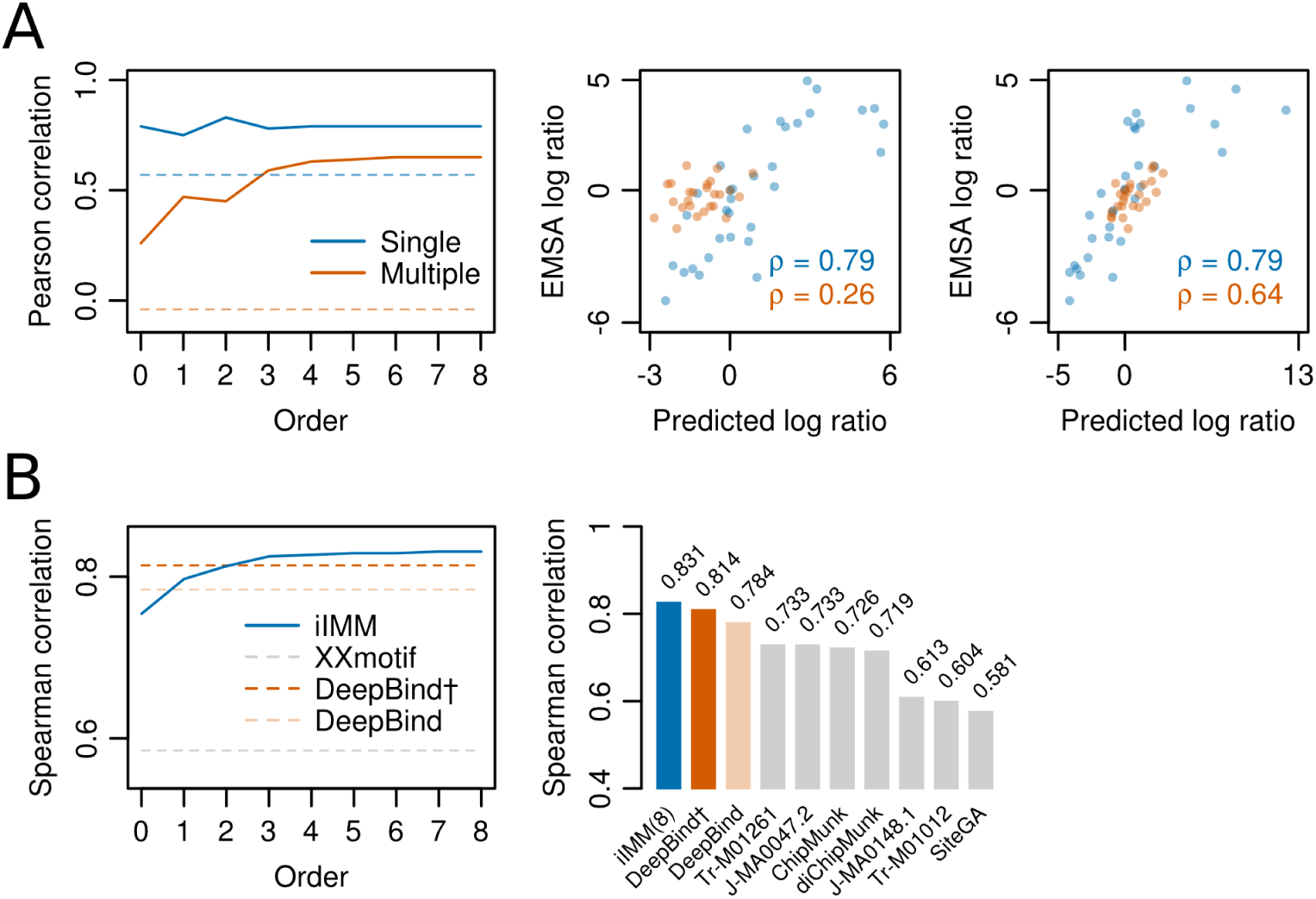
**Higher-order BMMs boost accuracy of binding affinity predictions for weak sites. (A)** Left: Pearson correlation between 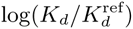 values for Klf4 binding to singly and multiply mutated consensus sites and 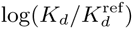 values predicted with models trained on Klf4-bound sequences from ChIP-seq. Solid lines: BMM models; dashed lines: PWMs from XXmotif. Middle: 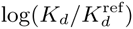 values measured by competitive EMSA assay versus values predicted by 0’th-order BMM trained on ChIP-seq data. Right: same but predictions from 5’th-order BMM. Affinities to weak, multiply mutated sites (orange) are very badly predicted using a PWM (correlation 0.26) but decently using a 5’th-order BMM (correlation 0.64). **(B)** Left: same as **A**, but showing Spearman rank correlations for FoxA2 binding affinities measured for 64 putative binding sites. DeepBind† and DeepBind differ only in model length (16 versus 24bp). Right: Spearman correlations of our 8’th-order BMM and various other methods, adopted from Alipanahi et al. [50].

To confirm these results, we performed a similar analysis on a dataset of competitive EMSA measurements for 64 double-stranded oligonucleotide probes containing potential FoxA2 binding sites [49]. These were correlated with predictions from the deep learning method DeepBind [50] and various other methods, which were trained on a FoxA2 ENCODE ChIP-seq dataset, and with prediction from a number of published PWMs for FoxA2. We repeated the analysis for our BMM predictions and obtained a Spearman correlation of *r* = 0.831, better than the best competing method, DeepBind (*r* = 0.814 and 0.784) (Figure 4B). This is remarkable, as no parameters were adjusted and our BMMs were not developed with the aim of quantitative prediction of binding affinities.

These results indicate that, at least for some factors, BMMs of order ≥ 3 are required to satisfactorily predict binding affinities to low-affinity binding sites. This is reflected in the information content of the higher-order sequence logos (Supplementary Figure S13).

### Predicting RNAP II transcription start sites

Whereas our previous analyses were based on simple motifs composed of a single binding site, we now assess higher-order Markov models for modelling complex motifs, regulatory regions that are targeted by multiple, cooperatively binding factors. Such complex motifs can be composed of multiple, non-obligatory submotifs with variable spacings and strengths.

The core promoter is the region of approximately ±50 bp around the transcription start site (TSS) that is required to initiate transcription. RNAP II core promoters can be classified into two classes [51], exhibiting a TSS distribution with a narrow peak (NP) or a broad peak (BP). In animals the former tend to be correlated with highly regulated genes and more frequently carry TATA-box and Initiator motifs, while the latter are correlated with housekeeping genes and have fewer, more poorly defined motifs.

We clustered and filtered TSSs measured in *D. melanogaster* by cap analysis of gene expression (CAGE) [52], resulting in 15 971 TSS clusters, assigned to 11 536 unique genes, which we classified into 7262 NP and 8709 BP core promoters. Furthermore, we modeled ribosomal protein (RP) gene core promoters using 92 core promoter sequences, corresponding to 86 unique RP genes listed in the RPG database [53].

We again used four-fold cross-validation, training on 75 % of the TSSs, testing on 25% of TSSs and pooling results of the four test sets. Because of the different peak widths for NP, BP and RP core promoters (90% of CAGE tags are contained in regions of 9bp, 47bp and 23bp around the peak mode, respectively), we took training sequences of lengths 109 bp, 147 bp, and 123bp around the mode of each TSS peak and trained models of length 101bp on them. We trained the 2’nd order background model on 501bp sequences centered around TSSs. For each test sequence of 501bp around a TSS, the position with the largest log-odds score was taken to be the predicted TSS. When the prediction was within 4bp, 23bp or 11bp in the case of NP, BP, and RP promoters, respectively, it counted as a true positive prediction, otherwise as false positive. The precision was the fraction of sequences with true positive predictions.

Figure 5A shows the positional distribution of predictions around NP TSSs for BMMs of order 0, 1 and 2, normalised to a random predictor with uniform density. While NP core promoters are relatively well predicted within ±4 bp by all three models, the 2’nd-order model achieves a precision of 76 %, 26 % higher than the precision of the PWM model (60 %). These improvements are reflected by a concomitant increase in total information content of the models (middle top). For even higher model orders, the precision saturates but, crucially, does not show any signs of overfitting (Supplementary Figure S16).

**Figure 5.**
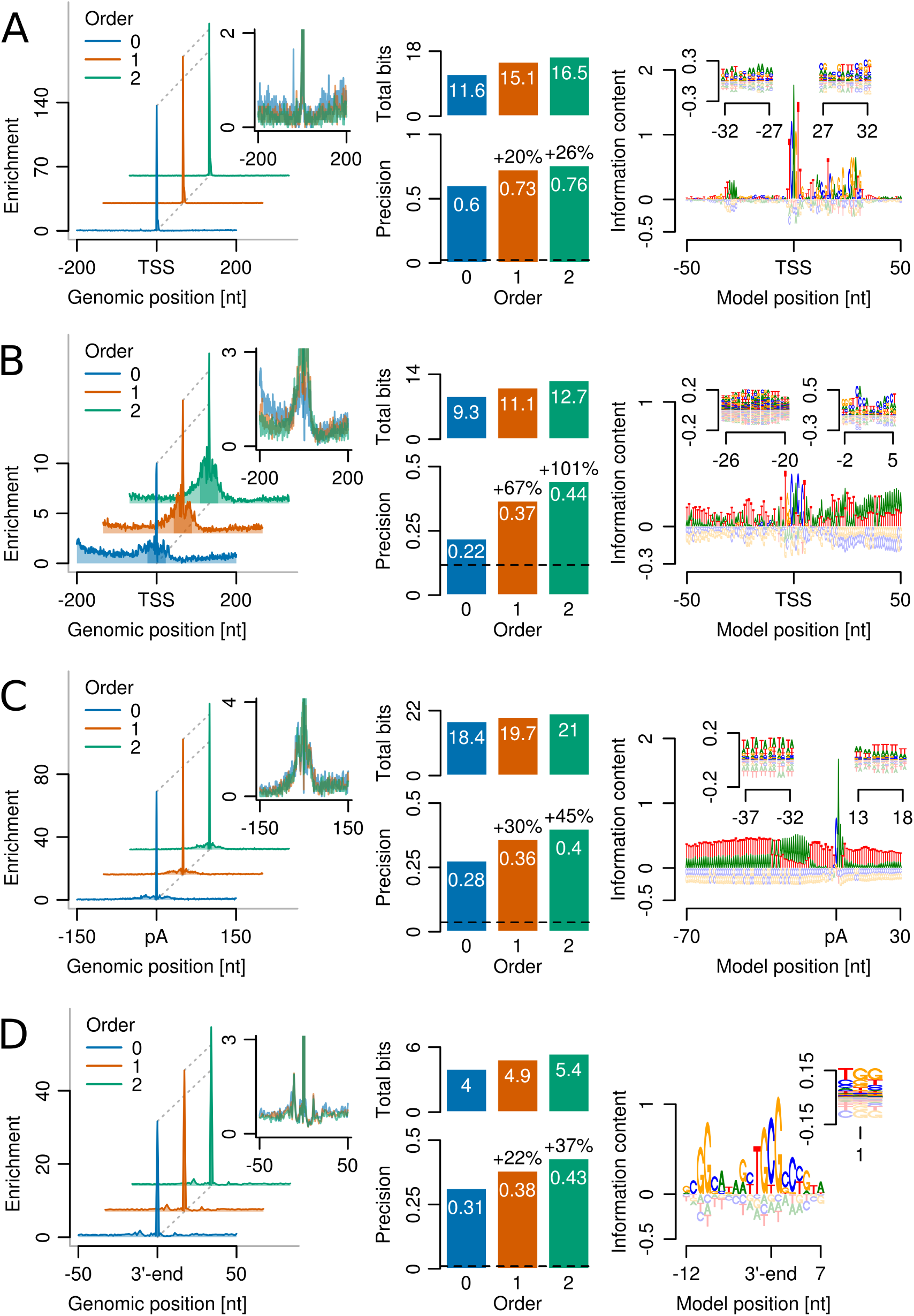
**Higher-order BMMs excel at predicting complex, multipartite motifs. (A)** Left: positional distribution of TSS predictions around measured TSSs of 7 262 narrow peak (NP) core promoters in *D. melanogaster*, normalised to random prediction with uniform density. Inset: same with expanded y-axis to show false predictions. Middle bottom: Fraction of sequences with correct predictions, defined to lie within 4bp of measured TSS peak mode. Dashed line: precision of random predictor. Middle top: Total information content in BMMs. Right: 0’th-order sequence logo of 2’nd-order BMM. Insets: 1’st-order sequence logos in region covering the TATA box (left) and the DPE and E-box motifs (right). **(B)** Same as **A** but for TSSs of 8709 broad peak (BP) promoters. Correct predictions are defined to lie within 23 bp of measured TSSs. Logo insets show 2’nd-and 1’st-order contributions. **(C)** Same as **A** but for polyadenylation (pA) sites from *S. cerevisiae*. Correct predictions are within 5bp of measured pA sites. Logo insets show 1’st-order contributions at efficiency and U-rich elements. (D) Same as A but for RNAP pause sites from *E. coli*. Correct predictions are within 0bp of measured pause sites.

The 0’th-order sequence logo of the 2’nd-order NP core promoter BMM (Figure 5A, right) reveals the Initiator motif at the TSS, the TATA box near -32 bp, and the motif ten and downstream promoter elements (MTE, DPE). The left inset shows 1’st-order dependencies in the region around the TATA box, which partly arise from the variable positioning of the TATA box with respect to the TSS. The right inset covers a region of overlapping DPE and E-box motifs and gives an idea how such overlapping, alternative motifs can be represented by a 1’st-order BMM. (See Supplementary Figure S14 for complete sequence logos.)

The BP TSSs are evidently much harder to predict than the NP TSSs (Figure 5B), owing to the scarcity and poor information content of their motifs. For difficult cases, however, BMMs show particularly clearcut gains: the precision achieved by a 2’nd-order BMM of 44 % is twice as high as for the 0’th-order model. Similar to NP core promoters, the BMM of BP core promoters represents sequence motifs that slightly vary in positioning and also distinct, overlapping motifs (Figure 5B, right). For instance, the Ohler6 element [54] and the DNA recognition element (DRE) are admixed but distinguishable in the 2’nd-order sequence logo (left inset). Similarly, the core promoter elements Ohler1 and Ohler7 [54] are overlapping each other but are distinguishable in higher orders (right inset).

The information density in higher orders of the NP and BP models seems rather low, but it sums up to considerable sizes across the modeled region (Supplementary Figures S14 and S15). We speculate that the nucleotide dependencies in our models reflect, at least in part, DNA structural properties of core promoter regions that contribute to TSS recognition [55].

Models of RP gene core promoter sequences do not profit from higher orders (Supplementary Figure S17A). However, despite the low number of sequence instances, even a 5’th-order BMM is not overfit (Supplementary Figure S18A).

### Prediction of polyadenylation sites

Sequence elements around the RNAP II polyadenylation (pA) site induce transcription termination by recruiting the cleavage and polyadenylation machinery to the pA site. In *S. cerevisiae*, 3′-end processing sequence signals were detected in the range from roughly 70 bp upstream to 30 bp downstream of the pA site [56] including the UA-rich efficiency element (EE), the A-rich positioning element (PE) and U-rich elements.

We extracted sequences of length 401 bp around 4 228 pA sites in *S. cerevisiae* from 4173 unique genes using major transcript isoform annotations [57]. The training and analysis was performed analagous to the core promoter prediction, using four-fold cross-validation. Training was done on the regions from — 70 bp to +30 bp around the pA sites and training of the BMM background model on the full 401 bp sequences. Testing was done on full-length 401 bp sequences, with true positive predictions defined to lie within 5bp of the annotated pA site.

Again, higher-order BMMs outperform PWM models by a wide margin, improving the 28% precision of the PWM model by 45% up to 40% for the 2’nd-order BMM (Figure 5C, left and middle bottom). The 2’nd-order BMM comprises all known 3′-end processing elements (Figure 5C, right). The 1’st-order correlations are necessary to model the EE (left inset) and the downstream T-rich region (right inset), which is revealed to be a poly(dA:dT) tract in higher orders (inset and Supplementary Figure S16). Again, even models of very high order did not suffer from overtraining (Supplementary Figure S18B).

### Prediction of bacterial RNAP pause sites

Pausing of RNAP during transcription has regulatory functions in RNA folding, recruitment of factors to mRNAs, and transcription termination. Larson *et al*. [58] measured RNAP pause sites in *E. coli* and *B. subtilis* using nascent elongating transcript sequencing (NET-seq) and identified 16 bp and 12 bp RNAP pause sequence signatures, respectively.

We extracted sequences of length 121 bp centered at 11 648 *E. coli* and 6 809 *B. subtilis* pause sites. 20bp models were trained on the regions from —12bp to +7bp around pause sites, which adds 2 bp in *E. coli* and 4 bp in *B. subtilis* to either end of the pause site motifs defined by Larson *et al*. The 2’nd order background model was trained on the entire genome. The assessment was analogous to the TSS and pA site predictions, using four-fold cross-validation and defining correct predictions as being precise within 0 bp.

The 0’th-order BMM predicts 31 % of *E. coli* pause sites correctly, the 1’st-order BMM increases the precision to 38 %, and the 2’nd-order BMMs to 43 % (Figure 5D). This suggests that pause sites might have a specific signature of DNA structural properties reflected in higher-order nucleotide dependencies. Off-site predictions up- and downstream of the pause index, e.g. at —11bp, are presumably caused by local similarities in the sequence features (Figure 5D, right).

Beside the GpG dinucleotide at the 5′-end of the RNA-DNA hybrid, 10 bp upstream of the 3′-end, another distinctive feature of the consensus sequence described by Larson *et al*. is the occurrence of TpG or CpG at the location of the 3′-end of the nascent transcript and incoming nucleoside triphospate. The CpG dinucleotide of the template strand was recently shown to inhibit elongation and induce G-to-A errors when spanning the active site of RNAP [59]. Our 2’nd-order BMM refines this signature by revealing that after a TpG a G is favoured, whereas CpG is more likely to be followed by a T or C (Figure 5D, right, inset).

In *B. subtilis*, the precision is only about half as high as in *E. coli*, but improvements through higher orders BMMs are more marked. The 3’rd-order BMM reaches 21 % precision, an increase of 55% over 0’th-order (Supplementary Figures S17B and S18C).

Pause site models differ substantially in all orders between the gram-negative *E. coli* and the gram-positive *B. subtilis*, except for a GpG dinucleotide at the upstream edge of the RNA-DNA hybrid and a pyrimidine at the downstream edge.

### Prediction of protein-RNA binding sites

In cells mRNAs are actively kept in a largely unfolded state by energy-dependent processes [60]. In contrast to DNA, which forms a relatively stiff double helix, mRNAs are therefore mostly single-stranded and extremely flexible. This leads to profound differences in the sequence specificity of RNA- versus DNA-binding. DNA sequences similar to the consensus sequence will usually be bound in a very similar overall protein-DNA conformation, whereas a single mutation from high-affinity RNA motif will usually cause the highly flexible mRNA to change its structure quite dramatically in order to minimise the binding energy. Such behaviour strongly violates the assumption that the binding energy can be approximated by independent energy contributions from each nucleotide, and it comes as no surprise that PWMs are poor models for RNA binding factors [61]. We were therefore wondering whether BMMs would be more appropriate.

We used a dataset of binding sites of 25 mRNP biogenesis factors from *S. cerevisiae* measured *in vivo* using PAR-CLIP [62, 63]. We extracted 25 nt sequences centered around the crosslinked uridines of the strongest 2 000 binding sites. We randomly sampled 20000 25 nt background sequences centered around uridines in the transcriptome of *S. cerevisiae*. Our analysis of the performance to discriminate between bound and background sequences proceeds in a way analogous to the benchmark in Figures 2 and 3.

Figure 6A shows the discriminative power of 2’nd-order BMMs compared to 0’th-order, PWM models. The performance of all models is low in comparison to DNA binders in Figure 2 when we consider that here we used a ratio of background to positive sequences of 10:1 instead of 100:1. BMMs outperformed PWM models for all RNA-binding proteins (*P* = 3 × 10^−8^, Wilcoxon one-sided signed-rank test, *n* = 25), in two cases doubling the pAUC values.

**Figure 6.**
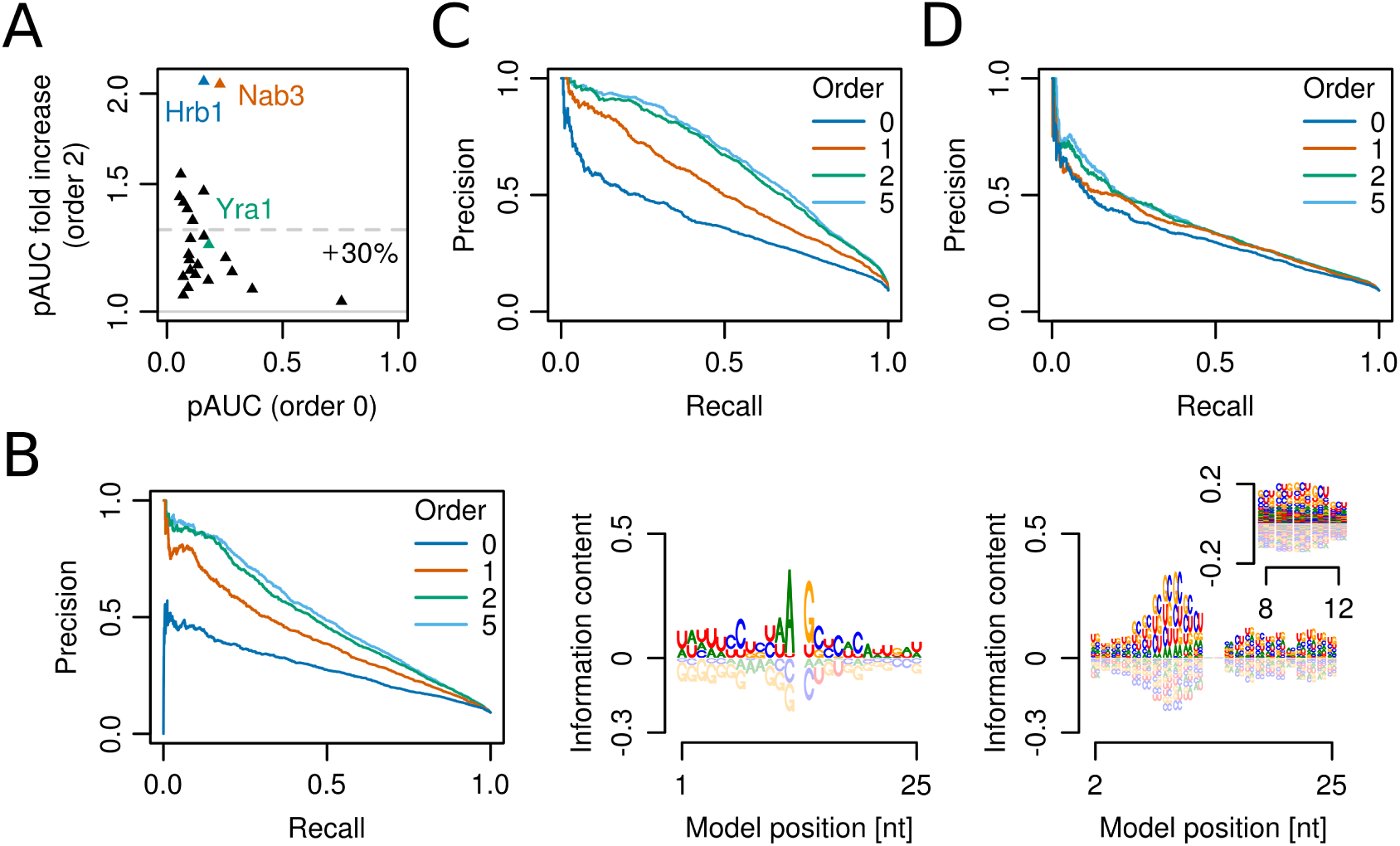
**Modelling the sequence specificity of RNA binding factors is challenging but improves with higher orders. (A)** Increase of prediction performance of 2’nd- versus 0’th-order BMMs for binding sites of 25 mRNP biogenesis factors from S. cerevisiae measured by PAR-CLIP. Dashed line: mean fold increase. **(B)** Left: Higher-order BMMs lead to sizeable gains in precision and recall for predicting Hrb1 binding sites. Right: sequence logos for order 0 and 1 of 2’nd-order BMM (central crosslinked U was removed from the 0’th-order logo). **(C,D)** Effect of BMM model order on prediction performance for Nab3 (C) and Yra1 (D).

The most drastic improvement was seen for the SR-like factor Hrb1 (Figure 6B). The crosslinked U at position 0 (not shown) is frequently flanked by A and G upstream and downstream, respectively (Figure 6, middle), which are also enriched around other RNA-binding protein crosslink sites (Supplementary Figure S19). In the 1’st- and 2’nd-order logos we can discern a CUG-rich region upstream of the crosslink site (Figure 6B, right), representing up to five successive CUG repeats, which cannot be learned by the PWM due to their variable positioning. Hrb1 was not known to bind CUG-rich sequences, but our observation makes sense in light of the fact that it contains three RRM domains just like CELF1, which is known to bind CUG-rich ssRNAs [64].

Nab3 profits greatly from higher orders (Figure 6C) because the Nab3 motifs UCUU and CUUG are learned together with the Nrd1 motifs UGUA and GUAG, which are enriched near Nab3 crosslink sites [65] (Supplementary Figure S19A). A typical, very moderate gain is seen for the Mex67 adaptor protein Yra1 (Supplementary Figure 6D and Figure S19B).

## Discussion

In this study we developed a Bayesian approach to train inhomogeneous Markov models that uses the conditional probabilities from lower order *k*-1 as prior for order *k*. The BMMs trained with this scheme can be regarded as a variation of interpolated Markov models. Unlike the various heuristic schemes that have been proposed for choosing the interpolation weights [34, 35, 36, 37], in our Bayesian approach we do not need to make any ad hoc choices and merely require two hyperparameters to set the strengths of the Bayesian priors for all model orders.

Our scheme also sidesteps the common approach of pruning the discrete dependency graph of the Markov model in order to limit the model’s complexity [23, 31, 32, 33]. Instead, we use continuous, soft, data-driven cut-offs which are effectively realised using Bayesian priors. We thereby avoid the cumbersome discrete optimisation of a dependency graph and can make use of simple and effective optimisation methods. This also allowed us to develop an EM-based algorithm for motif discovery using BMM training.

We tested our BMMs in a cross-validation setting on hundreds of ENCODE ChIP-seq datasets, RNAP II core promoter sequences and polyadenylation sites, bacterial RNAP pause sites and RNA-bound sites from PAR-CLIP measurements, using the same hyperparameters. On all datasets, BMMs yielded sizeable improvements, typically around 30% - 40% increase in precision. BMMs of order 5 led to a significant improvement (at significance level 0.0625) over PWMs on 97 % of the 446 ENCODE ChIP-seq datasets, while the performance was very similar on the remaining 3% (Figure 3C). For comparison, the Markov models recently introduced in the JASPAR database (TFFMs) showed significantly improved performance on only 21% of 96 ENCODE ChIP-seq datasets [30].

Given this success with modeling nucleotide correlations, why did k-mer-based methods not clearly outperform PWM-based models in the DREAM5 challenge [22]? First, whereas ChIP-seq measurements are done with full-length transcription factors, many of which possess multiple DNA-binding domains and binding partners, PBM measurements are mostly done with single DNA binding domains. Second, we think that, in contrast to ChIP-seq measurements, the amount of information present in a PBM measurement is often not enough to learn a model more detailed than a PWM. To see why this might be the case, we note that each 8-mer occurs 16 times and each 10-mer occurs once on the PBMs used in [22]. Due to the measurement noise, usually only the affinities to 8-mers are considered, since these can be averaged over the 16 measurements. However, for many DNA-binding domains the nucleotides flanking the core 8-mer probably have a considerable influence on binding strength. This means that the 16 measurements of each 8-mer are convoluted by the effects of the flanking nucleotides and could therefore be too unreliable to allow for training complex models with many parameters. Another way to look at the question is this: In a PBM measurement typically only 1% of probes, i.e. 400, are significantly bound and will carry most of the information, which might not be enough to estimate the (4^2^ - 1)×10 = 150 parameters of a 1’st-order model reliably enough.

Despite the theoretically high number of parameters at larger orders, our BMMs never deteriorated in performance with increasing order *k*, in contrast to simple inhomogeneous Markov models (Supplementary Figures S12, S18, and S20).

We developed higher-order sequence logos to visualise the information learned in the various orders on top of what is contained in lower orders. As illustrated in several examples, the logos could often explain the origin of the added value in higher orders.

For transcription factors, the added value is owed to variable submotif spacings, variable dimerization partners, and DNA shape readout, neither of which can be adequately learned with PWMs. Variable submotif spacings could also be learned by profile HMMs or mixtures of PWMs, however they lack flexibility to model dinucleotide preferences due to DNA shape readout and to describe more complex architectures with variable presence of motifs. Also, HMM training is slow as it requires running a forward-backward algorithm in each iteration of the EM algorithm.

Extension of the core transcription factor binding motif by 8bp improved 5’th-order BMMs by 19% but PWMs by only 5%, due to the ability of higher orders to model the DNA shape constraints around the core binding site. Implicitly learning structural properties might work better than explicitly including them in the model [66], since *any* DNA physical property will be reflected in specific, learnable oligonucleotide preferences.

We tested the ability of PWMs and BMMs trained on ChIP-seq data to predict binding affinities for two transcription factors measured by EMSA on datasets of sequences near their consensus binding sequences. BMMs improved predictions of PWMs and a number of other methods. The improvements are likely owed to weak sites more than a single substitution away from the consensus. On singly mutated sequences, the PWM predicted binding affinities as well as the BMM, while on doubly mutated sites it showed a dismal correlation of 0.26 while the BMM achieved 0.64 (Figure 4). Hence PWMs learn to predict mostly the high-affinity sites correctly, as the energies of all sequences a single mutation away from consensus can still be described with their simple energy model, and the breakdown of the PWM performance for doubly mutated sites reflects the breakdown of the additivity assumption according to which nucleotides contribute individually to the binding energy. As low-affinity sites have been reported to be important for the specificity and robustness of gene expression [67], improvements in predicting binding to weak sites will be important for quantitative modeling of transcriptional regulation.

Complex, multipartite motifs profited from the flexibility of BMMs to represent multiple submotifs at variable spacings and strengths. This flexibility may be useful to predict binding sites of cooperatively binding transcription factors, which often prefer certain spacings and orientations [15, 68]. It might prove particularly powerful for predicting binding sites of factors with multiple DNA-binding domains, such as ZnF transcription factors. ZnFs comprise up to a third of human transcription factors [69] and they contain on average 10 DNA-binding domains, which might partly explain why clearly defined motifs could be found for 8 % of them [3]. The striking prediction performance and specificity we observed for Znf143 (Supplementary Figure S9) and a 67-bp-long model of CTCF (Supplementary Figure S11) indicates that the complex binding sites of ZnF transcription factors with their multiple DNA-binding domains might be well predictable using BMMs.

At present, a limitation of BMMs in comparison to Bayesian network models (e.g. [32, 23]) is that nucleotide dependencies are only modeled within consecutive *k*-mers. Overcoming this would be useful for transcription factors that change their binding mode depending on the sequence and to learn complex motif architectures with correlated submotif occurrences, for example. It seems straightforward to generalise the presented approach by making each position *j* dependent on up to *k* not necessarily neighbouring upstream positions using a heuristic selection strategy. Because the described EM algorithm is fast – a 5’th-order BMM with 21 positions from 5000 sequences of 205bp is learned within 83s on a single core of a 3.4 GHz Intel Core i7-2600 CPU – we may choose a generous value *k*=5 or even higher in practice.

To further improve the performance of our BMM-based motif discovery tool BaMM!motif, we will learn automatically from the data (1) the hyperparameters and (2) the model length, a nontrivial task since the changing number of parameters precludes a simple optimisation of the likelihood or posterior probability. We will also (3) develop an efficient method to estimate the biological significance of discovered motifs, and (4) introduce positional priors. (5) Since BaMM!motif’s speed is at present limited by the seed motif discovery code from XXmotif, we will accelerate this code substantially. One future challenge will also be (6) to develop a rigorous approach that can deal with quantitative data such as fluorescence intensities from HT-SELEX and protein binding microarrays and peak strengths of ChIP-seq measurements.

## Conclusion

We have developed a Bayesian approach to train higher-order Markov models, which automatically adapts model complexity to the amount of available data position- and k-mer-specifically. The BMMs learned with this scheme were never overtrained, even at high orders. We developed the to our knowledge first method for learning the dependency graph among motif positions of higher-order Markov models that does not require a-priori knowledge of motif locations and that can hence be applied to de-novo motif discovery.

The most remarkable result of this study is the consistency with which higher-order BMMs yielded solid improvements across various heterogeneous datasets without requiring parameter tuning on each dataset and without a single case of failure. We can therefore answer affirmatively the question of whether nucleotide correlations are significant in transcription factor binding sites and other regulatory regions.

These results argue in favour of making the transition from PWMs to BMMs as the standard model to describe protein-DNA binding affinities and to offer BMM models in databases for regulatory and binding site motifs.

## Abbreviations

BaMM!motif: Bayesian Markov Model motif discovery; BMM: Bayesian Markov model; iMM: Inhomogeneous Markov model; MM: Markov model; pAUC: Partial area under the ROC curve; PWM: Position weight matrix; ROC: Receiver operating characteristic.

## Acknowledgements

We would like to thank Holger Hartmann for help with issues related to XXmotif, Mark Heron and Phillipp Torkler for help with core promoter and PAR-CLIP data, respectively, and all members of the Söoding lab for fruitful discussions.

